# Switchgrass Root Cell Wall Composition and Anatomy Vary with Depth, Suggesting Approaches for Trait Enhancement

**DOI:** 10.64898/2026.08.14.744798

**Authors:** Rahele Panahabadi, Jeremy B. Jewell, Ajaya Kumar Biswal, Nancy L. Engle, Niharika Nonavinakere Chandrakanth, Janel Poisson, Sushree S. Mohanty, Timothy J. Tschaplinski, Debra Mohnen, Anne E. Harman-Ware, Laura E. Bartley

## Abstract

Plant root cellular architecture and cell wall composition influence plant productivity, stress resilience, biotic interactions, and potentially soil carbon accumulation. This study establishes comprehensive compositional parameters for roots of a lowland switchgrass genotype, DVR3. Root traits were analyzed in 12.5 cm depth segments, from Zone 1 near the surface to Zone 4 down to 50 cm. Mean abundance (µg/mg) for major cell wall components included cellulose 470 ± 20, xylose 250 ± 20, lignin 170 ± 15, and total suberin 35 ± 5. Composition and cellular anatomy varied with depth, in a partially coordinated manner. Cross sections showed extensive aerenchyma in mature root regions despite greater root mass density, corresponding to abundant lignin and cellulose. Deep roots were enriched for pectin-associated traits, including arabinogalactan II, homogalacturonan, and arabinose-associated linkages. Suberin content did not vary significantly, though Casparian strip formation, endoderm and exoderm thickening, and suberin surface staining progressed with development. Similar trends in root lignin and specific root length were observed for another lowland switchgrass genotype, AP13. These results suggest that it may be possible to genetically enhance native switchgrass root chemistry to promote soil penetration and below-ground carbon accumulation by reducing variability with development, potentially via cell-type specific adjustments.

**Highlight:** Older, shallower switchgrass crown roots are enriched in lignin and cellulose, and deeper, younger roots are pectin-rich with juvenile cellular anatomy. A more uniform compositional distribution might enhance below-ground traits.

## Graphical Abstract_Alt: Alt text

Three column schematic summary of switchgrass root anatomy and composition across four 12.5-cm depth zones of a 50-cm root system. The first column is an image of the root zones prior to washing. Zone 1 represents older, shallow root segments and Zone 4 includes younger root segments down to the root tips. The second column is of representative cross-sections from each zone showing greater aerenchyma development in older root segments than in segments near the tip. The last column is a heatmap of major compositional features, showing higher cellulose, lignin, and xylose in Zone 1, higher pectin and nitrogen in Zone 4, and relatively little variation in suberin across zones.

## Introduction

Switchgrass (*Panicum virgatum* L.) is a leading bioeconomy crop because it produces high biomass while supporting ecosystem services, including soil carbon (C) accumulation that improves soil structure (Bates *et al*., 2022; Kibet *et al*., 2016; Morris *et al*., 2017; Schmer, 2008; Blanco-Canqui *et al*., 2014; Liebig *et al*., 2016; Kelly Slatten *et al*., 2023; Giabardo *et al*., 2026). For example, soil organic carbon (SOC) increased with five years of switchgrass growth by an average of 5.4 Mg C ha^-1^ at 0-30 cm across ten sites (Liebig *et al.,* 2008). Increasing SOC in agricultural systems may be an important strategy for carbon management, with global sequestration potential estimated at ∼4 Gt C year ¹ (Bossio *et al*., 2020). In addition, SOC improves nutrient availability, water retention, and soil fertility (Loria and Lal, 2025). Switchgrass roots can extend beyond three meters (Weaver, 1968), although root mass decreases with depth (Griffiths *et al*., 2022). Understanding switchgrass root composition and anatomy variation with depth supports optimization of both biomass production and ecosystem services.

New SOC is deposited most actively in topsoil (Heckman *et al*., 2023), typically defined as the top 30 cm below the surface. Under most conditions, root-, rather than shoot-, derived carbon preferentially contributes to stable forms of SOC (Pett-Ridge *et al*., 2021; Sokol *et al*., 2019a). The current paradigm posits that the largest source of root carbon is microbially processed root exudates, broadly defined (Cotrufo *et al*., 2013). Especially when nitrogen is abundant, labile compounds such as soluble sugars and non-cellulosic polysaccharides stimulate microbial activity, aggregate formation, and abundance of low molecular weight compounds that stably associate with soil minerals (Poirier *et al*., 2018a; Fierer and Schimel, 2003; Sokol *et al*., 2019b). More recalcitrant compounds, such as phenolic lignin and phenolic-aliphatic suberin, are often mineralized all the way to CO_2_ when microbes are abundant (Poirier *et al*., 2018b; Chen *et al*., 2017; Quigley and Kravchenko, 2022; Marschner *et al*., 2008; Cotrufo *et al*., 2013). Nonetheless, plant root detritus breakdown products processed by extracellular enzymes also adsorb to minerals, especially when microbial populations are low (Sokol *et al*., 2024) such as at greater depths or under water restriction (Sokol *et al*., 2026). Consistent with this, subsoils with reduced microbial activity harbor the most stable carbon (Gill *et al*., 1999; Li *et al*., 2023).

Although aboveground biomass composition has been extensively studied, root composition remains poorly characterized. Primary cell walls permit cellular elongation whereas late-forming secondary walls are usually thicker and less extensible. In both wall stages, cellulose provides structural support, while pectins contribute to porosity, hydration, and cell adhesion, especially in primary walls (Palin and Geitmann, 2012; Blamey *et al*., 2003; Lionetti *et al*., 2007). Lignin and suberin reinforce and create hydrophobic barriers in secondary walls (Vanholme *et al*., 2010; Saito and Fukushima, 2005; Baxter *et al*., 2009; Harman-Ware *et al*., 2021). Other matrix polysaccharides, especially xylan in grasses, form networks with cellulose and lignin (Scheller and Ulvskov, 2010; Tryfona *et al*., 2023). In grasses, hydroxycinnamic acids (HCAs), including esters of ferulic and *p*-coumaric acid, contribute to wall crosslinking and plant–microbe interactions (Chandrakanth *et al*., 2023; Bhattacharya *et al*., 2010; Gutjahr *et al.,* 2015).

Chemical features are integrated within specialized root tissues. The epidermis and root hairs enhance water and nutrient uptake, while suberin and lignin in the endodermis Casparian strip and exodermis regulate radial transport and protect against environmental stress (Steudle, 2000; Barberon, 2017; Geldner, 2013; Schreiber and Franke, 2011). Grass cortical cells frequently undergo cell death to produce aerenchyma, air-filled lacunae associated with improved gas conductance and reduced metabolic cost (Evans, 2004). Lignin of the sclerenchyma that underlies the exodermis enhances soil penetration (Steudle, 2000, Schneider *et al*. 2021).

Depth influences root function and resource acquisition. Phosphorus, potassium, and micronutrients are concentrated in upper soils, whereas water and nitrogen often move deeper in the soil profile (Lynch and Wojciechowski, 2015; Poirier *et al*., 2018a and 2018b). In grasses, deep axial and large lateral roots primarily acquire water and nitrogen, while fine lateral roots dominate nutrient uptake in topsoil (Lynch and Wojciechowski, 2015; Roumet *et al*., 2008).

Towards developing strategies for improving root traits, here, we analyzed root cell wall composition and anatomy across four developmental zones within the top 50 cm of a lowland switchgrass genotype, integrating anatomical observations with compositional profiling and glycosyl linkage analysis.

## Material and Methods

### Plant Material

The lowland switchgrass genotype Devils River 3 (DVR3; J210.A in Lovell *et al*., 2021) was obtained from T. Juenger and clonally propagated in the greenhouse for several years. For this experiment, a ramet initially consisting of a root crown and stems approximately 12 cm in diameter was grown for four months in 20 × 40 cm pots containing a 1:4 (v/v) mixture of sand and Sunshine Mix #3, daily irrigation, fertilization twice weekly with 20-10-20 fertilizer (200 ppm N), natural lighting with greenhouse temperature setpoints ranging from 20.0 to 23.9°C. The roots were harvested in November. The root mass was manually washed, and the total root mass in the pot was divided vertically into four parts, representing four biological replicates (Fig. 1A). Then each part was divided horizontally into four zones so that the root tips were included in zone 4. For consistency due to loss during washing and to focus on the bulk of the root biomass, lateral roots longer than 5 cm were removed. Then, each biological replicate was divided into two parts. One part was dried for measuring cell wall components, and the other part was preserved in 50% alcohol for microscopy. When analyzed separately, root tip sections were collected from within 0.5 mm of the root tip. From each zone, before alcohol-insoluble residue (AIR) preparation and de-starching (raw samples), some samples were dried and saved for suberin, nitrogen, and carbon analysis.

**Fig. 1.**
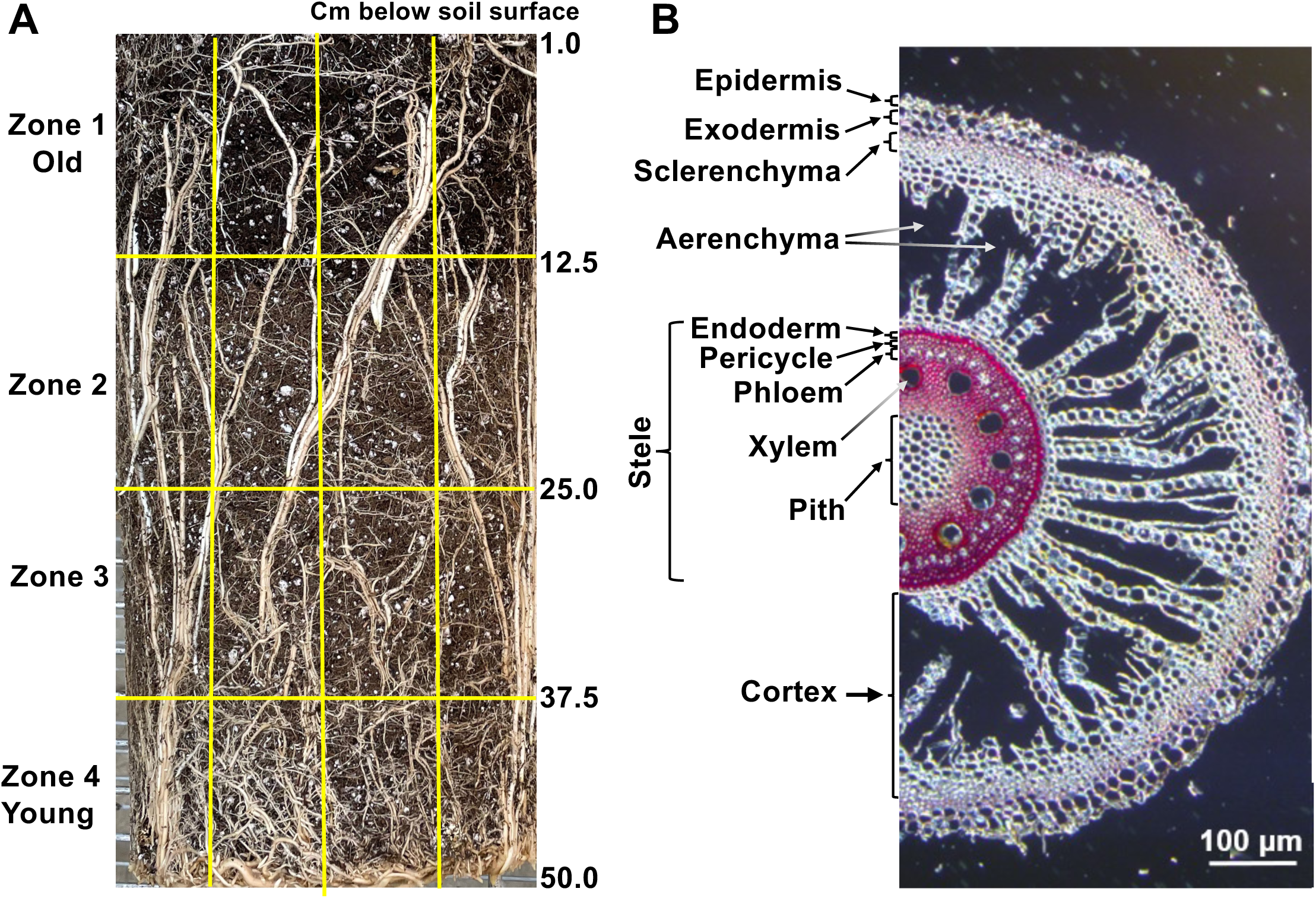
Zones of characterized switchgrass roots and root cellular anatomy. A) The switchgrass root system was divided into four ∼12.5 cm zones spanning older surface root segments (Zone 1) to young root tips (bottom of Zone 4). Root segments from each zone were separated into four longitudinal replicates, with long lateral roots (>5 cm) removed. Samples were split for microscopy and cell wall compositional analysis. B) A mature root cross section from Zone 1 displays characteristic cellular anatomy including phloroglucinol-HCl staining (fuchsia) of lignin in the exodermis, endodermis, and around the xylem and phloem. **Alt text_Fig. 1:** Two-panel figure illustrating the experimental sampling strategy and root anatomy in switchgrass. Panel A shows a photograph of the fibrous switchgrass root system divided into four 12.5 cm developmental zones from older surface roots (Zone 1) to younger root tips (Zone 4), with each zone separated into longitudinal replicates for microscopy and cell wall composition analyses. Panel B shows a part of a cross-section of a mature root stained with phloroglucinol-HCl, highlighting lignified tissues in the exodermis, endodermis, xylem, and phloem. Other cell layers are labeled. The cortex region is dominated by large spaces without cells, called aerenchyma.

In a related experiment, a clone of the switchgrass genotype AP13 (also identified as J206.A in Lovell *et al*., 2021) was split into three ramets of ∼5 cm diameter and each ramet was grown in a mesocosm (150 cm X 15 cm) filled with a mixture of sand, perlite, and vermiculite (12:7:1 by volume) from October to December, 2022, with light supplementation to 12 hours. Mesocosms were watered daily and fertilized as above 3 times weekly. After two months the mesocosms were emptied and plant roots segmented in 30 cm lengths, washed, stored in 50% ethanol at 4 °C until root tissue density and cell wall preparations were conducted.

### Root tissue density

Root tissue density (an approximation of, 1/root specific length) was calculated as the ratio between root length and dry root mass (cm•mg^−1^). Ten pieces of root for each Zone were chosen randomly and the mass and primary root length measured (without accounting for short lateral roots), followed by drying for 3 days at 45 °C. After drying, the mass was measured again, and density was estimated. The fresh weight to dry weight ratio of Zone 1 was 1.8:1 and for Zone 4 was 3:1.

### Sectioning, Staining and Microscopy

Healthy, undamaged roots of representative thickness were selected for sectioning from among the roots preserved in 50% ethanol per each Zone. Sections were prepared in three biological replicates from the top of each root zone (i.e., close to the soil surface) using a rapid-tome hand-sectioning device (Thomas *et al*., 2023). Root tip sections were prepared from the terminal 0.5 cm of the root. Bright-field images were collected using a Leica DMi3000 B inverted microscope (Leica Microsystems, Wetzlar, Germany). Lignin was stained with freshly prepared phloroglucinol–HCl, consisting of one part of concentrated HCl [37% (w/w)] and two parts of 3% (w/v) phloroglucinol in ethanol, with immediate mounting and examination via bright-field microscopy (Singh *et al.,* 2019). Pectin and lignin were simultaneously visualized via staining in 0.02% (w/v) Toluidine Blue O solution in distilled water for 5 min at room temperature followed by washing with distilled water to remove excess dye and immediate imaging (O’Brien *et al*., 1964). Suberin was stained with 0.03% Fluorol Yellow 088 (w/v) in lactic acid as previously described (Marhavý and Siddique, 2021). Briefly, the root was divided in half longitudinally and the cut surface laid flat on a slide and incubated with the Fluorol Yellow solution at 70 °C for 20 minutes, followed by washing in water for 1 minute at room temperature, counterstaining with 0.5% (w/v) aniline blue solution in water for 20 minutes at room temperature in the dark, and washing in water for 10 minutes. After staining, the cut surface was imaged using a Leica DMi3000 B inverted microscope (Leica Microsystems, Wetzlar, Germany) equipped with a GFP filter set.

### Root Biomass Preparation

The root segments for composition measurements were dried at 45 °C for three days. Dried samples were ground using a Wiley knife mill and passed through a 60-mesh sieve before further analysis.

### Carbon/Nitrogen Measurements

Carbon and nitrogen content of unextracted root samples was determined at the Stable Isotope Core Laboratory at Washington State University using a single pooled sample for each zone. Approximately 1.5 to 2 mg of dried and finely ground sample was encapsulated in tin, combusted to generate N and CO gases with an elemental analyzer (ECS 4010, Costech Analytical, Valencia, CA, USA), and separated using a continuous-flow gas chromatograph coupled to an isotope ratio mass spectrometer (Delta PlusXP, Thermo Finnigan, Bremen, Germany). Carbon and nitrogen contents were quantified using multi-point glutamic acid and acetanilid calibration following the laboratory’s standard protocols. Reported variation is estimated based on known instrumental precision.

### Preparation of Cell Wall Residues

For cell wall analysis, ground tissue (∼1 g) was destarched and extracted with alcohol to produce alcohol-insoluble residue (dsAIR) as previously described (Decker *et al*., 2015). Briefly, samples were transferred into tea bags and incubated in 0.1 M sodium acetate buffer (pH 5.0) containing glucoamylase (Rhizopus; TCI America, ∼1,600 AGU L ¹) and α-amylase (Bacillus; Sigma-Aldrich, 1.5% (v/v)) for 24 h at 55 °C with shaking at 120 rpm. Samples were then washed three times with distilled water and Soxhlet-extracted in ethanol for 24 h.

### Lignin Quantification Using Acetyl bromide

Lignin was quantified by the acetyl bromide method using a 96-well microplate. Briefly, 2 mg of dsAIR was incubated with 100 µL of freshly prepared acetyl bromide solution [25% (v/v) in acetic acid] in 2.0-mL screw-cap microcentrifuge tubes (Thermo Scientific, Cat. No. 3463) at 50 °C for 3 h in a thermomixer set to 1,050 rpm, with vortexing every 15 min during the final hour. Following incubation, 400 µL of 2 M NaOH and 70 µL of freshly prepared 0.5 M hydroxylamine hydrochloride were added. The mixture was vortexed, acidified with glacial acetic acid, and centrifuged at 12,000 rpm for 5 min. A 200 µL aliquot of the supernatant was transferred to a UV-compatible 96-well plate. Absorbance at 280 nm was used with the average extinction coefficient for grass aerial tissue (17.75 L g ¹ cm ¹) (Saito and Fukushima, 2005) and a path length of 0.55 cm using the Beer-Lambert equation. The use of an aerial tissue extinction coefficient might introduce errors in absolute quantification for root material but is expected to be reliable for relative quantification among related tissues.

### Lignin Monomer Determination by Pyrolysis Molecular Beam Mass Spectrometry

Py-MBMS analysis was performed using approximately 4 mg of root dsAIR. A Frontier PY2020 unit pyrolyzed samples at 500 °C for 30 s in 80 µL deactivated stainless steel cups. An Extrel Super-Sonic MBMS Model Max 1000 was used to collect mass spectral data from *m/z* 30 to 450 at 17 eV. Data were processed using Merlin Automation (V3) and R. Lignin-derived pyrolysates produce ions at *m/z* 120, 124 (G), 137 (G), 138 (G), 150 (G), 152, 154 (S), 164 (G), 167 (S), 168 (S), 178 (G), 180, 181, 182 (S), 194 (S), 208 (S) and 210 (S) where G indicates guaiacyl-derived ions, S indicates syringyl-derived ions, and other ions either derive from other lignin monomers or multiple sources. Ratios of S and G lignin monomer units (S/G) were obtained by dividing the sum of S-based ions by the sum of G-based ions using mean-normalized ion intensities.

### Analysis of Hydroxycinnamic Acids

Hydroxycinnamic acids (HCAs), ferulic acid (FA) and *p*-coumaric acid (*p*CA), were extracted from dsAIR into hemicellulosic and non-hemicellulosic fractions, the latter of which is enriched for lignin and suberin in roots. Briefly, 5.0 ± 0.5 mg of dsAIR was weighed into 2 mL screw-cap tubes and incubated with 0.5 mL of 0.05 M trifluoroacetic acid (TFA) with *trans*-cinnamic acid (*t*CA; 10 µL of a 1 mg mL ¹ solution) as an internal standard. Samples were incubated at 100 °C for five hours, cooled to room temperature, and centrifuged at maximum speed for five min. Then, the supernatant was carefully collected (approximately 450 µL), and the remaining pellet was washed with 50% (v/v) ethanol to remove residual soluble components. Subsequently, HCAs were released from both fractions by alkaline hydrolysis. All samples were incubated at 25 °C for exactly 24 h in a thermomixer and protected from light to prevent the photodegradation of ferulic acid (FA). Samples were then acidified to approximately pH 2 by the addition of 160 µL of concentrated hydrochloric acid [37% (w/w), ∼12 M], centrifuged for 5 min at 4000 rpm, and at least 250 µL of the clarified supernatant was transferred to glass vials and diluted 1:1 with water. High-performance liquid chromatography (HPLC) analysis was performed using a Shimadzu Nexera-i LC-2040C 3D Plus system, configured with LC-2060 modules and equipped with a photodiode array (PDA) detector. Samples were separated on a reverse-phase C18 column (Synergi™ 4 µm Fusion-RP 80 Å, 250 × 2.0 mm; Phenomenex) at a flow rate of 0.3 mL min ¹. The mobile phases consisted of solvent A (0.2% [v/v] trifluoroacetic acid in water) and solvent C (acetonitrile). The gradient program was as follows 14% C from 0 to 20 min, increased to 100% C from 20 to 25 min, held at 100% C until 33 min, returned to 14% C by 33 min, and held at 14% C until 50 min for column re-equilibration. The column temperature was maintained at 30 °C, and detection was carried out at 320 nm. Quantification was by peak area relative to external standards of *p*-coumaric acid and ferulic acid prepared in water at concentrations of 0.001, 0.003, 0.01, and 0.03 mg mL ¹.

### Monosaccharide Composition

In the main text, cell wall monosaccharides are reported as the average of measurements taken via high performance anion exchange chromatography with pulsed-amperometric detection (HPEAC-PAD) and trimethylsilyl (TMS) derivatization followed by gas chromatography mass spectrometry (GC-MS) detection approaches. Despite generally high agreement between the two methods, HPEAC consistently measured lower values for xylose than TMS, potentially due to incomplete hydrolysis by trifluoro acetate (TFA). The supplement includes the two separate measurements. For HPEAC-PAD analysis, similar to the method in Bartley *et al*. (2013), 5 mg of dsAIR was incubated in 2 mL tubes (Thermo Scientific Screw Cap Micro Tube, LOT number: 22350194) with 0.5 mL of 0.39 M TFA for 2 h at 120 °C with shaking and centrifugation every 30 min. After cooling to room temperature, samples were centrifuged at 12 k RPM for 10 min before transferring the supernatant into a new 2 mL-tube. The pellet was washed twice with cold 100% ethanol, and the washes were added to the TFA prior to drying via lyophilization. The resulting residue was resuspended in 0.5 mL of water and incubated with shaking at 1000 rpm at 70°C for 2 hours. After centrifugation of the resuspended sugars, 100 µL was removed and diluted in 900 µL of water. Monosaccharides were resolved using a Dionex ICS 6000 HPAEC-PAD operated by using Chromeleon software version 6.80 (ThermoFisher) using a Dionex CarboPac PA1 column. Monosaccharide standards were purchased from Sigma-Aldrich. The mobile phases consisted of water (solvent A), 0.1 M sodium hydroxide (solvent B), and 1 M sodium hydroxide (solvent C). Ion chromatography was performed at a flow rate of 0.4 mL min ¹. Prior to sample injection, the column was equilibrated for 15 min with 99% A and 1% B. The gradient program was as follows: 0-23.0 min, 99% A and 1% B; 23.1-41.0 min, 55% A and 45% C; and 41.01-45.0 min, 96% A and 4% C for column re-equilibration before the end of the run.

For trimethylsilyl (TMS) gas chromatography-mass spectrometry (GC-MS) analysis, as described by (York *et al*., 1985), 2 mg of dsAIR was hydrolyzed for 18 h at 80 °C in 400 μL 1 M methanolic–HCl and then derivatized with 300 μL of TriSil via heating to 80 °C for 20 min. The samples were resuspended in 150 μL hexane and 1 μL of sample was injected into the GC–MS for detection and quantification of sugars.

### Linkage analysis by GC-MS

Glycosyl linkage analysis was conducted via combined GC-MS of partially methylated alditol acetates as previously described (Black *et al.,* 2023). Briefly, samples were acetylated using N-methylimidazole and acetic anhydride in the ionic liquid 1-ethyl-3-methylimidazolium acetate. After dialyzing, the samples were methylated with potassium methylsulfinylmethylide and subsequently reduced with lithium aluminum deuteride in tetrahydrofuran. Permethylation was achieved by two rounds of treatment with sodium hydroxide and methyl iodide in dry dimethyl sulfoxide (Ciucanu and Kerek, 1984). The permethylated material was hydrolyzed with 2 M TFA for 2 hours in sealed tubes at 121 °C, followed by reduction with sodium borodeuteride and acetylation with acetic anhydride/TFA (York, 1985; Biswal *et al.,* 2015). The resulting PMAAs were analyzed using an Agilent 7890A gas chromatograph (Heiss, 2009) coupled to a 5975C mass selective detector in electron impact ionization mode. Chromatographic separation was performed on a 30 m Supelco SP-2331 bonded phase fused silica capillary column.

### Cellulose quantification

Cellulose content was determined on the pellet remaining after TFA hydrolysis of dsAIR for monosaccharide analyses as described (Bartley *et al*., 2013). Briefly, after TFA hydrolysate removal and washing of the pellet with ethanol, 175 µl 72% H SO was added to the pellets. Then, tubes were capped, vortexed, and incubated for 45 min at room temperature with mixing at 500 rpm. After 30 min, 1225 µl H O was added, samples were vortexed and then centrifuged for 5 min at max speed. Five µl of supernatant was mixed with 45 µl of H O for both samples and standards (0, 0.125, 0.25, 0.5, 0.75, 1, and 2 mg ml ¹ glucose in H O). Fresh anthrone reagent (100 µl; 2 mg anthrone ml ¹ in 98% H SO) was then added to each 50 µl sample or standard, followed by incubation at 80 °C for 30 min and cooling to room temperature. Thereafter, 120 µl was transferred to a clear 96-well plate and absorbance was measured at 625 nm, and cellulose concentration was estimated based on relative response to external standards.

### Suberin Monomer Analysis by GC-MS

Suberin preparations were carried out based on Jenkin and Molina (2015). Tissue de-lipidation was conducted at room temperature on a rotisserie rotator and all solvents contained 0.01% (w/v) butylated hydroxytoluene (BHT). All procedures for suberin analysis were performed in glass tubes with polytetrafluoroethylene-faced screw cap lids. Dried and milled root samples (∼50 mg) were extracted once with isopropanol for 2 hr and a second time overnight. Next, pellets were extracted with 2:1 CHCl_3_:MeOH (v/v) overnight, then overnight in 2:1 MeOH:CHCl_3_ (v/v).

Tubes were spun at 800 rcf for 10 min after each extraction, and the supernatant discarded. De-lipidated pellets were allowed to dry uncapped overnight in a chemical fume hood, then dried for 3 days under vacuum in a glass desiccator containing anhydrous desiccant (DRIERITE).

For depolymerization, a reaction mix containing 0.9 mL methyl acetate, 1.5 mL sodium methoxide (25% in methanol), 3.6 mL methanol and 25 µg methyl heptadecanoate (internal standard) was added to each tube. Sample tubes were capped tightly, vortexed, and placed in a 60 °C bath for 2 hr, with vortexing every 15 min. After cooling, 1.5 mL glacial acetic acid and 10 mL dichloromethane were added to each tube, then 0.5 M NaCl was added to fill each tube. Tubes were vortexed, centrifuged at 800 rcf for 10 min, then the lower organic layer was collected to a new tube. This was repeated, and both extractions pooled. The extract was washed twice with 0.5 M NaCl, discarding the upper aqueous layer. To remove remaining water, a small amount of anhydrous sodium sulfate was added to each tube and vortexed. The next day, tubes were centrifuged at 800 rcf for 10 min and the organic phase was collected to a new pre-weighed tube. Solution was evaporated to dryness under a gentle stream of nitrogen and tubes were sealed.

For GC-MS, the dried residue was dissolved in 2 mL 80% ethanol, then 1 mL was transferred to a new tube and dried again under a nitrogen stream. Acetonitrile (500 µL) and N-Methyl-N-(trimethylsilyl) trifluoroacetamide (500 µL) containing 1% trimethylchlorosilane was added to the dried residue and the sample was heated at 70 °C for an hour to generate trimethylsilyl derivatives. One µL of each sample was injected 2 days later to an Agilent 7890A gas chromatograph coupled to a 5975C inert XL mass spectrometer with the GC operating in splitless mode with the MS in electron impact ionization mode. Carrier gas flow (helium) was 1 mL/min and scanning was set for 40-650 amu. Chromatographic separation was performed with a Restek Rtx-5MS with Integra-Guard column (30 m x 0.25mm ID x 0.25µm df). Compounds were identified utilizing a Wiley Registry/NIST Mass Spectral Library in addition to a large user-created spectral database. Suberin monomer quantities are reported as methyl heptadecanoate equivalents.

### Statistical Analysis

One-way ANOVA was performed to evaluate differences among root zones, followed by Tukey’s honestly significant difference (HSD) test for multiple comparisons using the agricolae package in R (version 4.2.0). Differences were considered statistically significant at P < 0.05. Simple linear regression were used to evaluate the agreement between HPAEC and TMS measurements and scatter plots, regression lines, and 1:1 reference lines were generated in R using the ggplot2 package. Heatmaps were generated in R using the ggplot2, tidyr, and dplyr packages.

## Results

### Cellular anatomy and density

Fifty centimeter, pot-grown roots from the lowland switchgrass genotype, DVR3, were washed and divided into four 12.5 cm-zones to examine developmental and depth-associated anatomical and compositional variation below the plant crown (Fig. 1A). Zones were numbered from the soil surface downward, with Zone 1 representing the oldest root tissue near the crown and Zone 4 representing the youngest tissue, including actively growing root tips. Fig. 1B provides an overview of switchgrass root cellular organization.

Histochemical stains and microscopy revealed developmental differences in root cellular anatomy and composition across the root zones (Fig. 2). Cross-sections from Zones 1 and 2 contained extensive cortical aerenchyma, largely absent in Zone 4. Nonetheless, younger, deeper root zones had lower tissue density (Table 1, Supplemental Figure 1) and suggests that density is not driven by aerenchyma inabundance but increased secondary wall deposition. Consistent with this, phloroglucinol staining, which detects lignification, was more intense in Zones 1 and 2 than in younger root zones (Fig. 2A and 2B) and was especially evident in the stele and exodermal tissues. Endodermal lignin staining, indicating formation of the Casparian strip, was absent in root tips, but well formed at the top of Zone 4.

**Fig. 2.**
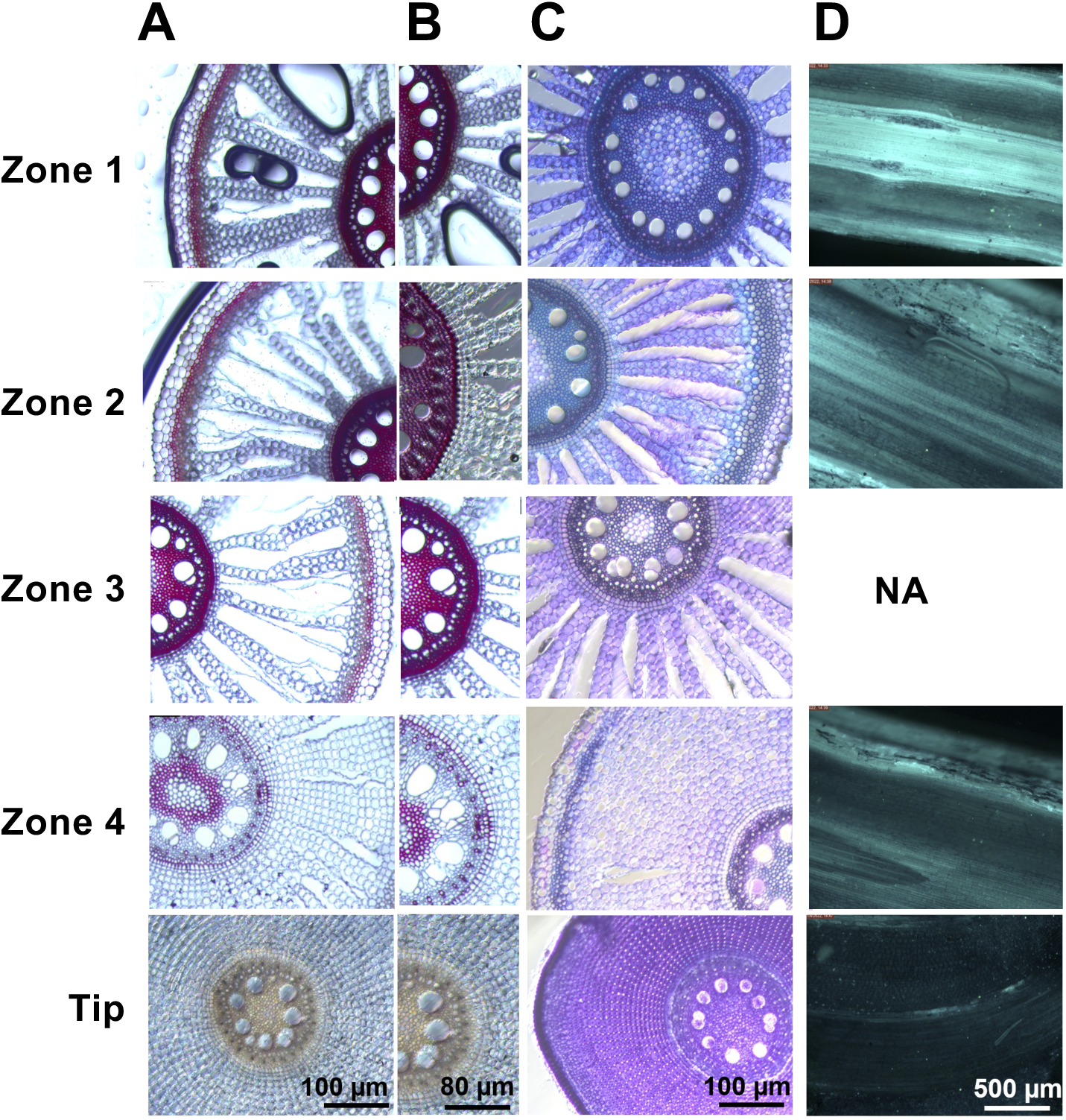
Switchgrass root anatomy and cell wall composition vary across developmental zones.. A-C) Cross-sections from different root zones and primary root tips show anatomical changes and shifts in cell wall composition during development. A-B) Phloroglucinol staining (fuchsia) revealed progressive lignin accumulation in walls of specific cells. B) Image magnification highlights endoderm cell development. C) Toluidine blue staining differentiated lignified (blue) and pectin-rich (purple) cell walls. D) Fluorol Yellow 088 staining of root longitudinal sections (cyan) showed increasing suberin deposition from the root tip to upper root zones, with little or no signal at the tip and strong staining in mature zones. See Fig. 1B for cell type labels. **Alt text_Fig. 2**: Four-panel figure showing anatomical and cell wall changes across switchgrass root developmental zones. Cross-sections reveal increasing tissue differentiation and progressive lignification as roots mature, with enlarged views highlighting endodermal development. Toluidine blue staining distinguishes lignified and pectin-rich cell walls, while Fluorol Yellow 088 staining shows little suberin at the root tip and progressively stronger suberin deposition in more mature root regions.

**Table 1.** Density and cell wall composition of switchgrass root zones. Values are presented as means ± standard deviation, with N = 4 biological replicates for cell wall composition and N = 3 biological replicates for suberin data. For each component, values that do not share letters are significantly different via ANOVA (P < 0.05) followed by Tukey’s HSD. Lateral roots longer than ∼5 cm were removed. Monosaccharide data represent the average of the HPAEC and TMS methods. Major suberin components are included here; all suberin data are provided in Table S2. Figure S1 displays these results in heatmap form.

| Depth (cm) | 0 - 12.5 | 12.5 - 25 | 25 - 37.5 | 37.5 - 50 |
| --- | --- | --- | --- | --- |
| Components | Zone 1 | Zone 2 | Zone 3 | Zone 4 <sup>†</sup> |
| Root tissue density (mg/cm) | 15 ± 2 <sup>a</sup> | 12 ± 1 <sup>ab</sup> | 11 ± 2 <sup>b</sup> | 9 ± 1 <sup>b</sup> |
| Carbon (μg/mg) | 470 ± 10 | 470 ± 10 | 460 ± 10 | 460 ± 10 |
| Nitrogen (μg/mg) | 8 ± 1 | 8 ± 1 | 11 ± 1 | 14 ± 1 |
| Carbon:Nitrogen | 57 ± 1 | 60 ± 1 | 41 ± 1 | 32 ± 1 |
| Cell Wall Total (μg/mg) <sup>^</sup> | 1090 ± 30 <sup>a</sup> | 1000 ± 30 <sup>b</sup> | 960 ± 30 <sup>b</sup> | 960 ± 20 <sup>b</sup> |
| Cellulose (μg/mg) | 520 ± 10 <sup>a</sup> | 460 ± 10 <sup>b</sup> | 450 ± 10 <sup>b</sup> | 440 ± 15 <sup>b</sup> |
| Lignin (μg/mg) | 190 ± 10 <sup>a</sup> | 170 ± 10 <sup>b</sup> | 160 ± 10 <sup>b</sup> | 150 ± 10 <sup>b</sup> |
| pCA-pellet (μg/mg) | 6.7 ± 0.2 <sup>b</sup> | 7.0 ± 0.1 <sup>ab</sup> | 7.8 ± 0.5 <sup>a</sup> | 7.2 ± 0.2 <sup>ab</sup> |
| FA-pellet (μg/mg) | 2.3 ± 0.1 <sup>a</sup> | 2.6 ± 0.1 <sup>a</sup> | 2.7 ± 0.2 <sup>a</sup> | 2.6 ± 0.2 <sup>a</sup> |
| S/G ratio | 0.48 ± 0.01 <sup>b</sup> | 0.50 ± 0.01 <sup>ab</sup> | 0.48 ± 0.01 <sup>b</sup> | 0.51 ± 0.01 <sup>a</sup> |
| Sum of matrix monosaccharides (μg/mg) | 370 ± 30 <sup>a</sup> | 360 ± 20 <sup>ab</sup> | 340 ± 20 <sup>ab</sup> | 340 ± 5 <sup>b</sup> |
| Xylose (μg/mg) | 270 ± 30 <sup>a</sup> | 260 ± 25 <sup>a</sup> | 230 ± 20 <sup>b</sup> | 225 ± 5 <sup>b</sup> |
| Arabinose (μg/mg) | 40 ± 2 <sup>bc</sup> | 36 ± 2 <sup>c</sup> | 42 ± 3 <sup>ab</sup> | 45 ± 1 <sup>a</sup> |
| pCA-xylan (μg/mg) | 3.2 ± 0.1 <sup>a</sup> | 3.4 ± 0.2 <sup>a</sup> | 3.2 ± 0.1 <sup>a</sup> | 3.4 ± 0.2 <sup>a</sup> |
| FA-xylan (μg/mg) | 2.6 ± 0.1 <sup>a</sup> | 3.0 ± 0.1 <sup>ab</sup> | 3.1 ± 0.1 <sup>ab</sup> | 3.2 ± 0.3 <sup>b</sup> |
| Glucuronic acid (μg/mg) | 0.8 ± 0.1 <sup>b</sup> | 1.0 ± 0.2 <sup>ab</sup> | 1.5 ± 0.3 <sup>a</sup> | 1.6 ± 0.2 <sup>a</sup> |
| Arabinose/Xylose | 0.15 ± 0.01 <sup>b</sup> | 0.15 ± 0.01 <sup>b</sup> | 0.18 ± 0.02 <sup>a</sup> | 0.20 ± 0.01 <sup>a</sup> |
| Arabinose/FA_Xylan | 15.0 ± 0.5 <sup>a</sup> | 12.0 ± 0.5 <sup>c</sup> | 13.5 ± 0.5 <sup>bc</sup> | 14.0 ± 0.5 <sup>ab</sup> |
| Glucose (μg/mg) | 32 ± 1 <sup>b</sup> | 32 ± 1 <sup>b</sup> | 37 ± 1 <sup>ab</sup> | 41 ± 2 <sup>a</sup> |
| Galactose (μg/mg) | 14 ± 1 <sup>b</sup> | 14 ± 1 <sup>b</sup> | 15 ± 1 <sup>ab</sup> | 16 ± 1 <sup>a</sup> |
| Pectin (GalA+Rha; μg/mg) | 4 ± 1 <sup>b</sup> | 4 ± 1 <sup>b</sup> | 6 ± 1 <sup>a</sup> | 8 ± 1 <sup>a</sup> |
| Mannose (μg/mg) | 3.5 ± 0.5 <sup>a</sup> | 3.5 ± 0.5 <sup>a</sup> | 3.5 ± 0.5 <sup>a</sup> | 3.5 ± 0.5 <sup>a</sup> |
| Galacturonic acid (μg/mg) | 3.0 ± 0.5 <sup>b</sup> | 3.0 ± 0.5 <sup>b</sup> | 3.5 ± 0.5 <sup>ab</sup> | 4.5 ± 0.5 <sup>a</sup> |
| Rhamnose (μg/mg) | 2.0 ± 0.2 <sup>b</sup> | 1.5 ± 0.2 <sup>b</sup> | 3.0 ± 0.5 <sup>a</sup> | 3.5 ± 0.5 <sup>a</sup> |
| Fucose (μg/mg) | 0.5 ± 0.05 <sup>b</sup> | 0.5 ± 0.05 <sup>b</sup> | 0.7 ± 0.05 <sup>a</sup> | 0.8 ± 0.05 <sup>a</sup> |
| Rhamnose/GalA | 0.7 ± 0.1 <sup>a</sup> | 0.6 ± 0.08 <sup>a</sup> | 0.9 ± 0.02 <sup>a</sup> | 0.7 ± 0.07 <sup>a</sup> |
| Sum of suberin components without HCA (μg/mg) | 9.5 ± 0.3 <sup>a</sup> | 8.5 ± 0.2 <sup>a</sup> | 9.1 ± 0.7 <sup>a</sup> | 8.6 ± 0.5 <sup>a</sup> |
| 16-OH_16:0 (μg/mg) | 2.2 ± 0.5 <sup>a</sup> | 1.6 ± 0.1 <sup>a</sup> | 1.8 ± 0.3 <sup>a</sup> | 1.9 ± 0.3 <sup>a</sup> |
| Vanillin (μg/mg) | 1.2 ± 0.2 <sup>a</sup> | 1.1 ± 0.1 <sup>ab</sup> | 1.1 ± 0.1 <sup>ab</sup> | 0.8 ± 0.1 <sup>b</sup> |
| 4-hydroxybenzaldehyde (μg/mg) | 1.0 ± 0.2 <sup>a</sup> | 0.9 ± 0.1 <sup>ab</sup> | 1.0 ± 0.1 <sup>a</sup> | 0.6 ± 0.1 <sup>b</sup> |
| 18-OH_18:1 (μg/mg) | 1.1 ± 0.2 <sup>a</sup> | 0.9 ± 0.1 <sup>a</sup> | 1.0 ± 0.2 <sup>a</sup> | 1.1 ± 0.1 <sup>a</sup> |

Toluidine blue staining corroborated contrasting developmental patterns. Younger roots and root tips exhibited strong purple staining within cortical tissues, indicating abundance of pectin-rich primary walls and consistent with active cell expansion in developing tissues (O’Brien *et al*., 1964; Mori *et al*., 1996).

Fluorol Yellow staining, which binds to aliphatic layers of suberin, further demonstrated developmental differences among root zones. Longitudinal sections of Zone 1 roots exhibited strong fluorescence, whereas root tips gave little signal (Fig. 2D). These observations agree with work in other species showing progressive suberization during root maturation (Ursache *et al*., 2021; Cantó-Pastor *et al*., 2024). Together, the anatomical observations indicated a developmental transition from pectin-rich expanding tissues in younger, deeper roots toward lignified and suberized tissues in mature roots closer to the crown.

### Carbon and nitrogen

Carbon and nitrogen analyses of total root biomass showed that carbon content remained relatively constant across all developmental zones at 45-47% dry weight (Table 1, Supplementary Figure S1). In contrast, nitrogen per unit mass was substantially greater in younger tissues, ranging from 1.4% in Zone 4 to 0.8% in Zone 1, consistent with greater metabolic activity and lower secondary wall deposition in actively growing roots.

### Variation in polysaccharide and lignin composition along development

We examined composition and abundance of cell wall sugars, glycosyl linkage, lignin, hydroxycinnamates, and suberin-associated compounds (Table 1, Supplementary Fig. S1). As expected, younger, deeper root segments (Zones 3 and 4) were enriched in traits associated with primary walls and actively growing tissues, whereas older root segments near the crown (Zones 1 and 2) were enriched in secondary wall-associated traits (Table 1).

Switchgrass root cell walls contained 40-50% cellulose, 34-37% matrix polysaccharides, and 15-20% lignin. Hydroxycinnamates represented about 1.6% of the wall, with *p*-Coumaric acid (*p*CA) associated with the acid-resistant pellet fraction (*p*CA-pellet), which in roots includes lignin- and suberin-associated material, representing 0.6-0.7% of cell wall residue. Acid-soluble *p*CA associated with xylan (*p*CA-xylan) represented ∼0.3-0.4%, while ferulic acid associated with xylan (FA-xylan) accounted for ∼0.2-0.3%.

Most components varied significantly with depth and development (1-way ANOVA, P < 0.05), though variation was not perfectly coordinated. Cellulose and lignin were both most abundant in Zone 1 and progressively decreased toward the root tip. Cellulose content declined 15% from 520 µg/mg in Zone 1 to 440 µg/mg in Zone 4. Lignin decreased 21%, from 190 µg/mg in Zone 1 to 150 µg/mg in Zone 4. Trends were consistent with phloroglucinol staining patterns. In contrast, *p*CA associated with the acid-resistant pellet fraction showed the lowest abundance in Zone 1 and the greatest abundance in Zone 3. Dissociation between *p*CA abundance and lignin accumulation has been observed previously in grasses (Lin *et al*., 2016), which might be confounded by *p*CA incorporation into both lignin and suberin. Despite developmental differences in lignin abundance, the syringyl-to-guaiacyl (S/G) ratio remained 0.5 across all zones, suggesting no evidence of monolignol composition regulation at least among the relatively large root segments examined here.

Acid-solubilized monosaccharide content was determined by both sugar electrochemical detection (HPEAC-PAD) and GC-MS of trimethylsilyl (TMS) derivatives. Biswal *et al*. (2017) reported that these methods provide comparable quantitative results. Accordingly, monosaccharide abundance measured by HPAEC and TMS in the present study were strongly correlated (R = 0.98-0.99), though TMS sugars were consistently slightly higher than sugars by HPEAC-PAD (Supplementary Fig. S2). Hence, we report the monosaccharides as the average of the two methods. The complete dataset is available in on-line.

Monosaccharide composition and glycosyl linkage analyses revealed depth-dependent differences in matrix polysaccharide composition (Table 1; Table S1). Except for xylose, which was 23% more abundant in Zone 1 than in Zone 4 (270 vs. 225 µg mg ¹), most monosaccharides diminished as development proceeded, including arabinose, glucuronic acid, non-crystalline cellulose-derived glucose, and galactose, whereas mannose remained relatively constant across zones. These results are consistent with depletion (or dilution) of primary wall polysaccharides from glycoprotein-associated glycans and xylan substitutions as root cells age. Indeed, in the linkage analysis, the predominant heteroxylan backbone linkage, 4-linked xylose (4-Xyl*p*), was 16 mol% in Zone 4, but increased to 21 mol% in Zone 1, and 2,4-Xyl*p* and 3,4-Xylp linkage abundances were greater in Zones 3 and 4 than Zones 1 and 2 (Table S1), all consistent with greater substitution of heteroxylan, and a greater Ara/Xyl ratio, in younger tissues.

Monosaccharide and glycosyl linkage analysis also supported greater abundance of pectin and other minor cell wall matrix polysaccharides in younger roots. Pectin, calculated as the sum of galacturonic acid and rhamnose, was more abundant in Zone 4 compared to Zone 1 (0.8% vs. 0.4% dry weight). Glycosyl linkage analysis revealed greater abundance of 4-linked galacturonic acid (4-GalA*p*), terminal rhamnose (t-Rha*p*), and 2-linked rhamnose (2-Rha*p*) in younger/deeper roots (Table S1). These linkages are associated with homogalacturonan and rhamnogalacturonan I, supporting increased pectin abundance in actively growing tissues, also reflected in the strong toluidine blue staining of Zone 4 (Fig. 2C). Type II arabinogalactans, characterized by 3-, 6-, and 3,6-linked β-D-galactose residues (Pettolino *et al*., 2012; Pfeifer *et al*., 2020), were also enriched in Zones 3 and 4, relative to mature root segments. Increased abundance of terminal arabinofuranose (t-Ara*f*), 4-linked arabinopyranose or 5-linked arabinofuranose (4-Ara*p* or 5-Ara*f*), terminal rhamnose (t-Rha*p*), and terminal galactose (t-Gal*p*) further supported greater relative abundance of heteroxylan, arabinogalactan- and pectin-associated polymers in developing versus mature root segments.

### Suberin composition

GC-MS analysis of depolymerized root wall extracts revealed that the total abundance of putative suberin-associated monomers was 30 µg/mg root biomass, similar to values reported for other species (Baxter *et al*., 2009). However, the extracted monomer profile was dominated by hydroxycinnamic acids (HCAs), with coumaric acid and ferulic acid representing 42% and 33% of the total signal, respectively. Because HCAs are also major constituents of grass arabinoxylan and lignin (at ∼16 µg HCA/mg, Table 1), they cannot be attributed exclusively to suberin polymers, suggesting that when HCAs are included total suberin abundance may be overestimated, with a non-HCA suberin abundance of ∼9 µg/mg.

Fatty acids, hydroxy-fatty acids, and dioic acids with chain lengths from C16 to C28 collectively represented 17% (68% with HCAs excluded) of the total extracted monomer pool and most did not differ significantly among zones, though the 28:0 fatty acid was significantly lower in younger samples (Table S2). Additional compounds detected included vanillin, acetosyringone, hydroxybenzaldehyde, and sinapyl alcohol, collectively representing ∼9% (24% with HCAs excluded) of total signal, some of which were significantly reduced in Zone 4 relative to the mature zones. Overall, the suberin-associated polyester fraction showed relatively modest variation across the four developmental zones compared with polysaccharide and lignin-associated traits.

### Verification in another genotype

We examined root tissue density and lignin content in a tall pot system for another lowland switchgrass genotype, AP13. The results confirm the general trends reported above, with ∼14% more ABSL content in the top 30 cm than the bottom 30 cm, and ∼73% lower root density in young, deep roots compared to older ones (Supplementary Fig. S3).

## Discussion

Root structure and content development influence root mechanics, microbial interactions, decomposition dynamics, and long-term soil carbon persistence. In the transport roots of the switchgrass lowland genotypes characterized here, we observed developmental gradients in the anatomy and cell wall composition. Younger, deeper root segments were enriched in pectin- and arabinogalactan-associated polysaccharides, whereas mature root segments accumulated greater lignin, cellulose, and xylose. Though transport roots are the dominant root mass, especially in lowland switchgrass (Griffiths *et al*., 2022), removal of longer fine lateral roots likely reduced the detected abundance of pectin and other primary root components at shallower depths.

The root developmental progression was not completely coordinated and suggests control by distinct regulatory processes. Suberin content did not vary quantitatively, within the large segments characterized, though was absent from root tips; lignin and cellulose were significantly more abundant only in Zone 1 vs. Zones 2-4; pectic components and xylose showed a statistically significant transition between Zone 2 and Zone 3; aerenchyma developed between Zone 3 and Zone 4; and endodermal thickening occurred in the between 0.5 and 10 cm of the root tip. Depending on the compositional and structural targets, these contrasts can inform the experimental design and interpretation of future molecular profiling. For example, late-season root collection might reduce developmental differences among genotypes and simplify interpretation of shallow root compositional differences. On the other hand, among other properties of deep root tips, composition could be particularly important for soil penetration and carbon stability.

Hydroxycinnamic acids represent ∼1.6% of switchgrass root cell wall, with FA likely contributing to covalent cell wall polymer crosslinking (Chandrakanth *et al*., 2023), and *p*CA on xylan potentially reducing non-covalent cellulose-hemicellulose interactions (Pathare *et al*., 2024). Unlike aerial tissues in which the xylan-associated ratio of FA to *p*CA is ≥ 3:1 (Bartley *et al*., 2013; Tian *et al*., 2021), FA and *p*CA were equally abundant in the matrix polysaccharide fraction (HCA-xylan). This suggests that the main function (and related gene expression) for *p*CA-xylan may be in root processes. Of possible relevance, cell wall HCAs are produced in response to fungi (Gutjahr *et al*., 2015) and may modulate such interactions.

Root structure and content variation may influence root biophysical properties. Increased lignification contributes to root tensile strength and forms diffusion barriers inhibiting water and solute movement (Schneider, 2022; Reyt *et al*., 2021). The greater lignin abundance observed in the mature upper zone is consistent with lignin deposition increasing progressively as roots development (Zhu and Barros, 2025). Increasing lignin accumulation during early root maturation could further enhance switchgrass soil penetration and radial transport control. However, these capabilities would need to be balanced with continued elongation capacity that is likely supported by the observed greater abundance of pectin-associated monosaccharides and linkages, including galacturonic acid- and rhamnose-associated structures (Dash *et al*., 2023). Staining demonstrated progressive deposition of suberin with root maturation consistent with previous reports of suberization with development (Geldner, 2013; Andersen *et al*., 2018). However, GC-MS analysis detected only small differences in total depolymerized suberin-associated monomers across zones, suggesting that suberin endodermal and exodermal barriers are constructed soon after root cell differentiation and do not continue to accumulate.

Root content may influence SOC long-term stabilization (>100 years; Poirier *et al*., 2018b). Accumulation of plant carbon inputs in soil reflects a balance between occlusion from microbial processing versus metabolism through microbial mineralization to CO_2_ and/or loss through leaching (Fearnside and Barbosa, 1998; Cotrufo *et al*., 2013; Poirier *et al*., 2018b; Sokol *et al*., 2019; Angst *et al*., 2023). This balance is influenced by microbial activity, soil characteristics, and moisture, which among other factors are influenced by depth, with subsoils (∼>30 cm) having less microbial activity (Angst *et al*., 2023). While mineral-associated organic material (MAOM, < 53 µm) is typically the most stable of soil carbon components; particulate organic material (POM) can also be stable for thousands of years under certain conditions (Angst *et al*., 2023). MAOM production is thought to be favored by abundant sugary root exudates, but also high plant root detritus quality, including low lignin abundance, high C:N ratio, and high pectin abundance (Cotrufo *et al*., 2013; Poirier 2019b). Under conditions of relatively low microbial activity, MAOM can form with only minimal extracellular microbial processing of plant detritus (Sokol *et al*., 2019), retaining a stronger plant-derived chemical signature (Sokol *et al*., 2024; Rinehart *et al*., 2026). Thus, increasing the compositional uniformity of switchgrass roots across development could promote more readily digested root detritus higher in the soil column and more stable detritus deeper. More specifically, reducing lignin content in the stele and enhancing pectin content of cortex cells late in development could enhance MAOM, and vice versa early in development to enhance POM, respectively. In addition, the positive influence of S-lignin and *p*CA-derived phenolics on MAOM formation (Rinehart *et al*., 2026), suggests that genetically increasing these components throughout development might enhance MAOM.

## Conclusion

Establishing a framework for future studies, we present benchmark data suggesting that switchgrass roots exhibit partially coordinated developmental gradients in anatomy and cell wall composition across depth-associated zones. This opens opportunities for studying variation observed in other genotypes and genetic control of composition, expanding to plant-microbe interactions for synergistic enhancement of plant and soil properties, and probing relationships between these properties and nutrient use and recycling. We hypothesize that reducing the developmental variability of root composition by promoting lignin and *p*CA early in development and pectin later in development may enhance root-soil penetration, water and nutrient acquisition, and long-term soil carbon storage in many soil ecosystems. Thus, understanding and manipulating these developmental trajectories could provide new opportunities for breeding and engineering crops with optimized root traits that improve plant performance and increase ecosystem services for the bioeconomy.

## Supplementary Data

The following supplementary data are available at JXB online.

**Supplementary Figure S1.** Chemical variation across switchgrass root zones.

**Supplementary Figure S2.** Correlations between monosaccharide concentrations determined by HPAEC and TMS.

**Supplementary Table S1.** Glycosyl linkage analysis of switchgrass root zones.

**Supplementary Table S2.** Abundance of suberin monomer components across root zones.

## Acknowledgements

We thank members of the Center for Bioenergy Innovation and the Institute of Biological Chemistry for support and technical assistance.

## Author contributions

RP: conceptualization, methodology, investigation, formal analysis, data curation, visualization, writing – original draft; JBJ: methodology, investigation, formal analysis, writing – original draft, writing – review & editing; AKB: methodology, investigation, formal analysis, writing – review & editing; NE: investigation, formal analysis, writing – review & editing; NNC: methodology, writing – review & editing; JP: investigation, writing – review & editing; SSM: investigation; TT: methodology, writing – review & editing; AEH-W: funding acquisition, investigation, formal analysis, writing – review & editing; DM: conceptualization, writing – review & editing; LEB: conceptualization, supervision, project administration, funding acquisition, writing – original draft, writing – review & editing.

## Data Availability

The root images and complete datasets for root composition reported here are available at DOI: https://doi.org/10.5281/zenodo.21045502

## Abbreviations

AGI: arabinogalactan I
AGII: arabinogalactan II
AIR: alcohol-insoluble residue
Ara*f*: arabinofuranose
Ara*p*: arabinopyranose
C: carbon
C:N: carbon-to-nitrogen ratio
dsAIR: destarched alcohol-insoluble residue
FA: ferulic acid
GalA: galacturonic acid
GalA*p*: galacturonic acid in the pyranose form
Gal*p*: galactopyranose
Glc*p*: glucopyranose;HCA hydroxycinnamic acid
HG: homogalacturonan
HM: heteromannan
HX: heteroxylan
MAOM: mineral-associated organic matter
MLG: mixed-linkage glucan
*p*CA: *p*-coumaric acid
POM: particulate organic matter
RGI: rhamnogalacturonan I
RGII: rhamnogalacturonan II
Rha: rhamnose
Rha*p*: rhamnopyranose
S/G: syringyl-to-guaiacyl ratio
SOC: soil organic carbon
XG: xyloglucan
Xyl*p*: xylopyranose.

