## Supplemental Tables and Figures for "Switchgrass Root Cell Wall Composition and Anatomy Vary with Depth, Suggesting Approaches for Trait Enhancement"

**Table S1.** **Glycosyl linkage analysis of switchgrass root zones.** Values are expressed as mol% of total detected sugars. Linkages are grouped according to deduced polymer assignments (Pettolino *et al.*, 2012). Glycosyl linkage analysis was performed using pooled biological replicates for each root zone; therefore, statistical analysis was not performed. A dash indicates that the linkage was not detected.

|  | **Depth (cm)** | **0 - 12.5** | **12.5 - 25** | **25 – 37.5** | **37.5 - 50** |
| --- | --- | --- | --- | --- | --- |
| **Deduced linkage** | **Polysaccharide assignments** | **Zone 1** | **Zone 2** | **Zone 3** | **Zone 4** |
| t-Ara*f* | RGII/AGI/AGII/HX/RGI | 1.5 | 1.5 | 1.6 | 1.7 |
| t-Ara*p* | AGII | 0.1 | 0.1 | 0.1 | 0.2 |
| 3,4-Ara*p* or 3,5-Ara*f* | Arabinan | 0.1 | — | — | 0.2 |
| 3-Ara*f* | RGI | 0.8 | 0.5 | 0.8 | 0.9 |
|  | Total Arabinose | 1 | 0.6 | 0.9 | 1.3 |
| 4-Ara*p* or 5-Ara*f* | Arabinan/RGI | 0.6 | 0.4 | 0.7 | 0.7 |
| t-Rha*p* | RGII/AGII | — | 0.1 | 0.3 | 0.5 |
| 2-Rha*p* | RGI | 0.2 | 0.1 | 0.4 | 0.6 |
| t-Xyl*p* | XG/Pectin/other | 0.8 | 0.8 | 0.7 | 0.7 |
| 2-Xyl*p* | XG/RGII | 2.3 | 1.9 | 2.1 | 1.7 |
| 4-Xyl*p* | HX | 20.4 | 18.8 | 16 | 15.5 |
| 3,4-Xyl*p* | HX | 7.5 | 6.7 | 6.5 | 4.3 |
| 2,4-Xyl*p* | HX | 2 | 2.3 | 2 | 1.4 |
|  | Total HX | 29.9 | 27.8 | 24.5 | 21.2 |
| 4-Man*p* | HM | 0.2 | 0.2 | 0.2 | 0.2 |
| t-GalA*p* | HG/RGI/RGII | — | — | 0.1 | 0.1 |
| 4-GalA*p* | HG/RGI | 0.5 | 0.3 | 0.8 | 1.3 |
| t-Gal*p* | RGI/RGII/XG/HX/AGII | 2.3 | 1.4 | 2.4 | 2.8 |
| 3-Gal*p* | AGII/RGI | 0.7 | 0.5 | 0.9 | 0.9 |
| 3,6-Gal*p* | AGII/RGI | 0.3 | 0.3 | 0.3 | 0.4 |
| 6-Gal*p* | AGII/RGI/RGII | 0.2 | 0.2 | 0.1 | 0.2 |
|  | Total | 1.2 | 1 | 1.3 | 1.5 |
| 4-Gal*p* | AGI/RGI | 0.2 | 0.2 | 0.3 | 0.3 |
| 4-Glc*p* | XG/HM/Cellulose/other | 45.6 | 44.1 | 46.9 | 54.5 |
| 4,6-Glc*p* | XG/starch | 4.4 | 2.9 | 2 | 2.5 |
| t-Glc*p* | MLG/other | 1.5 | 7 | 7.5 | 2 |
| 2-Glc*p* | starch/other | — | 0.5 | 0.7 | — |
| 2,4-Glc*p* | starch/other | 2.2 | 2.4 | 1.4 | 0.8 |
| 3,4-Glc*p* | MLG/other | 1.9 | 1.6 | 1 | 1.1 |

**Table S2. Abundance of suberin monomer components in different zones of mature switchgrass roots.** Values are presented as means ± standard deviation (N = 3) and are expressed as µg mg⁻¹ heptadecanoic acid equivalents. HCA represents the sum of ferulic acid (FA) and *p*-coumaric acid (*p*CA). Different letters indicate significant differences as determined by Tukey's HSD post hoc test following one-way ANOVA (P < 0.05).

| **Depth (cm)** | **0 - 12.5** | **12.5 - 25** | **25 - 37.5** | **37.5 - 50** |
| --- | --- | --- | --- | --- |
| **Components** | **Zone 1** | **Zone 2** | **Zone 3** | **Zone 4** |
| *p*CA (µg/mg) | 15.7 ± 0.8 ^a^ | 15.3 ± 0.9 ^a^ | 17.0 ± 0.4 ^a^ | 13.4 ± 0.3 ^a^ |
| FA (µg/mg) | 12.4 ± 0.2 ^a^ | 11.6 ± 0.4 ^a^ | 13.0 ± 2 ^a^ | 11.8 ± 0.8 ^b^ |
| 16-OH_16:0 (µg/mg) | 2.2 ± 0.5 ^a^ | 1.7 ± 0.1 ^a^ | 1.8 ± 0.3 ^a^ | 1.9 ± 0.3 ^a^ |
| Vanillin (µg/mg) | 1.2 ± 0.2 ^a^ | 1.1 ± 0.1 ^ab^ | 1.1 ± 0.1 ^ab^ | 0.8 ± 0.1 ^b^ |
| 18-OH_18:1 (µg/mg) | 1.1 ± 0.2 ^a^ | 0.9 ± 0.1 ^a^ | 1.0 ± 0.2 ^a^ | 1.1 ± 0.1 ^a^ |
| 4-hydroxybenzaldehyde (µg/mg) | 1.0 ± 0.2 ^a^ | 0.9 ± 0.02 ^ab^ | 1.0 ± 0.2 ^a^ | 0.6 ± 0.07 ^b^ |
| 16:0 (µg/mg) | 0.6 ± 0.04 ^a^ | 0.6 ± 0.08 ^a^ | 0.8 ± 0.2^a^ | 0.7 ± 0.1 ^a^ |
| 18:0 (µg/mg) | 0.67 ± 0.06 ^a^ | 0.60 ± 0.05 ^a^ | 0.80 ± 0.02 ^a^ | 0.68 ± 0.01 ^a^ |
| C16_dioic_acid (µg/mg) | 0.58 ± 0.03 ^a^ | 0.47 ± 0.04 ^b^ | 0.51 ± 0.03 ^ab^ | 0.50 ± 0.05 ^ab^ |
| Acetosyringone (µg/mg) | 0.55 ± 0.04 ^ab^ | 0.65 ± 0.04 ^a^ | 0.60 ± 0.08 ^a^ | 0.50 ± 0.02 ^b^ |
| 16:1 (µg/mg) | 0.3 ± 0.2 ^b^ | 0.5 ± 0.1 ^a^ | 0.5 ± 0.2 ^a^ | 0.5 ± 0.1 ^a^ |
| 20:0_dioic_acid (µg/mg) | 0.17 ± 0.08 ^a^ | 0.25 ± 0.06 ^a^ | 0.25 ± 0.09 ^a^ | 0.30 ± 0.08 ^a^ |
| 18:1_dioic_acid (µg/mg) | 0.180 ± 0.009 ^a^ | 0.155 ± 0.005 ^a^ | 0.165 ± 0.009 ^a^ | 0.180 ± 0.02 ^a^ |
| Sinapyl_alcohol (µg/mg) | 0.10 ± 0.04 ^a^ | 0.15 ± 0.01 ^a^ | 0.13 ± 0.02 ^a^ | 0.10 ± 0.06 ^a^ |
| 20:0 (µg/mg) | 0.100 ± 0.002 ^a^ | 0.087 ± 0.002 ^b^ | 0.092 ± 0.004 ^ab^ | 0.094 ± 0.005 ^ab^ |
| 18:1 (µg/mg) | 0.05 ± 0.02 ^a^ | 0.04 ± 0.06 ^a^ | 0.04 ± 0.02 ^a^ | 0.05 ± 0.01 a |
| 18-OH_18:0 (µg/mg) | 0.041 ± 0.009 ^a^ | 0.040 ± 0.002 ^a^ | 0.040 ± 0.006 ^a^ | 0.035 ± 0.004 ^a^ |
| 22:0 (µg/mg) | 0.026 ± 0.001 ^ab^ | 0.022 ± 0.002 ^c^ | 0.022 ± 0.001 ^bc^ | 0.027 ± 0.002 ^a^ |
| 22-OH_22:0 (µg/mg) | 0.015 ± 0.035 ^ab^ | 0.010 ± 0.003 ^b^ | 0.013 ± 0.002 ^ab^ | 0.017 ± 0.025 ^a^ |
| 24:0 (µg/mg) | 0.071 ± 0.005 ^a^ | 0.060 ± 0.001 ^b^ | 0.055 ± 0.004 ^b^ | 0.070 ± 0.007 ^a^ |
| 24-OH_24:0 (µg/mg) | 0.05 ± 0.02 ^a^ | 0.04 ± 0.03 ^a^ | 0.05 ± 0.01 ^a^ | 0.065 ± 0.009 ^a^ |
| 26:0 (µg/mg) | 0.014 ± 0.001 ^a^ | 0.012 ± 0.003 ^a^ | 0.011 ± 0.01 ^a^ | 0.012 ± 0.010 ^a^ |
| 28:0 (µg/mg) | 0.050 ± 0.005 ^a^ | 0.044 ± 0.001 ^ab^ | 0.040 ± 0.002 ^bc^ | 0.033 ± 0.004 ^c^ |
| 26-OH_26:0 (µg/mg) | 0.01 ± 0.05 ^a^ | 0.01 ± 0.01 ^a^ | 0.01 ± 0.03 ^a^ | 0.02 ± 0.02 ^a^ |
| Sum suberin with HCA (µg/mg) | 37 ± 1 ^a^ | 35 ± 2 ^a^ | 37 ± 5 ^a^ | 34 ± 2 ^a^ |
| Sum suberin no HCAs (µg/mg) | 9.5 ± 0.3 ^a^ | 8.5 ± 0.2 ^a^ | 9.1 ± 0.7 ^a^ | 8.6 ± 0.6 ^a^ |

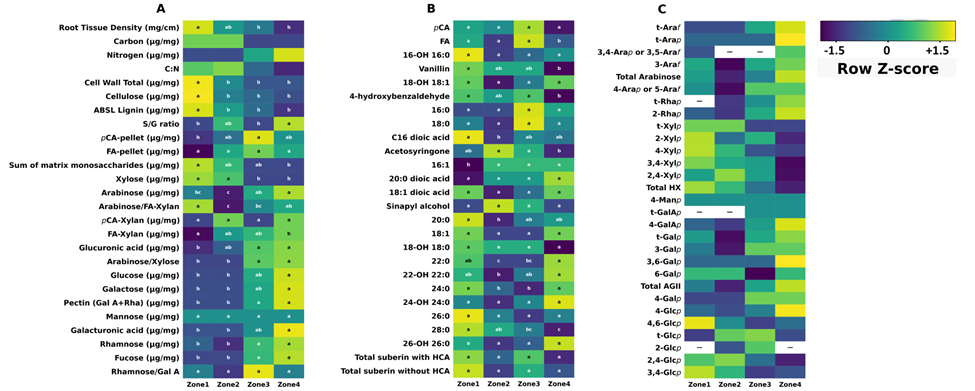
**Supplementary Figure S1.** Chemical variation across switchgrass root developmental zones. Values are row-scaled Z-scores, with blue indicating low abundance and yellow indicating high abundance. White cells containing a dash (—) indicate unavailable values, which were excluded from row-wise Z-score calculations. Zone 1 represents the upper 12.5 cm below the crown, whereas Zone 4 represents the deepest root region containing growing root tips. Lateral roots longer than ~5 cm were removed before analysis. (A) Structural cell wall traits, including lignin, cellulose, hydroxycinnamic acids, monosaccharides (average of HPAEC and TMS measurements), pectin-related traits, S/G ratio, and derived compositional ratios (see Table 1 for details). Cell wall total represents the sum of hydroxycinnamic acids, acetyl bromide-soluble lignin (ABSL lignin), and total monosaccharides. (B) Base-released soluble phenolics and suberin-derived aliphatic monomers (µg/g), including vanillin, acetosyringone, ω-hydroxy acids, and dicarboxylic acids. The suberin total represents the sum of quantified suberin lipid and aromatic components, without glycerol. (C) Glycosyl linkage analysis grouped by inferred polysaccharide assignments, including arabinan-, heteroxylan-, pectin-, and glucan-associated structures. Glycosyl linkage analysis was performed using pooled biological replicates; therefore, Tukey’s HSD post hoc test was not conducted. For all other panels, different letters indicate significant differences as determined by Tukey’s HSD post hoc test following ANOVA (P < 0.05). Values are in Table 1 and Table S1.

**Alt text_ Supplementary Figure S1**: Three columns of heatmaps showing row-normalized (Z-score) chemical variation across four switchgrass root developmental zones. Colors range from blue (lower relative abundance) to yellow (higher relative abundance), and white cells indicate unavailable values that were excluded from normalization. Panel A summarizes structural cell wall composition, Panel B summarizes soluble phenolics and suberin-associated compounds, and Panel C summarizes glycosyl linkage profiles grouped by inferred polysaccharides. The heatmaps show that multiple cell wall components change systematically with root developmental stage

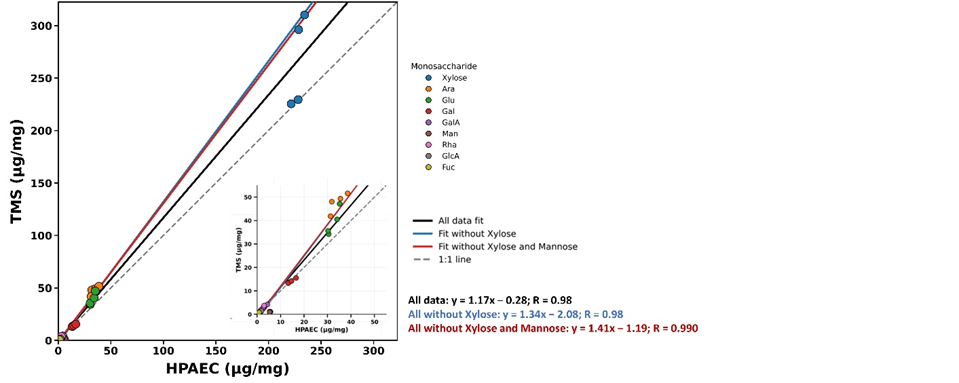

**Supplementary Figure S2:** Correlations between monosaccharide concentrations determined by HPAEC and TMS. The left panel shows the relationship between HPAEC and TMS measurements for all quantified monosaccharides. Because xylose is much more abundant than the other sugars, it dominates the overall distribution. The inset shows a magnified view of the lower concentration range to improve visualization of the remaining monosaccharides. Regression lines are shown for all monosaccharides, excluding xylose, and excluding both xylose and mannose.

**Alt text_ Supplementary Figure S2:** Scatter plot comparing monosaccharide concentrations measured by HPAEC and TMS. Most data points follow a positive linear relationship, indicating good agreement between the two analytical methods. Because xylose is much more abundant than the other monosaccharides, it dominates the main plot. An inset enlarges the lower concentration range to show the remaining sugars more clearly, with regression lines illustrating the relationships after excluding xylose and then excluding both xylose and mannose

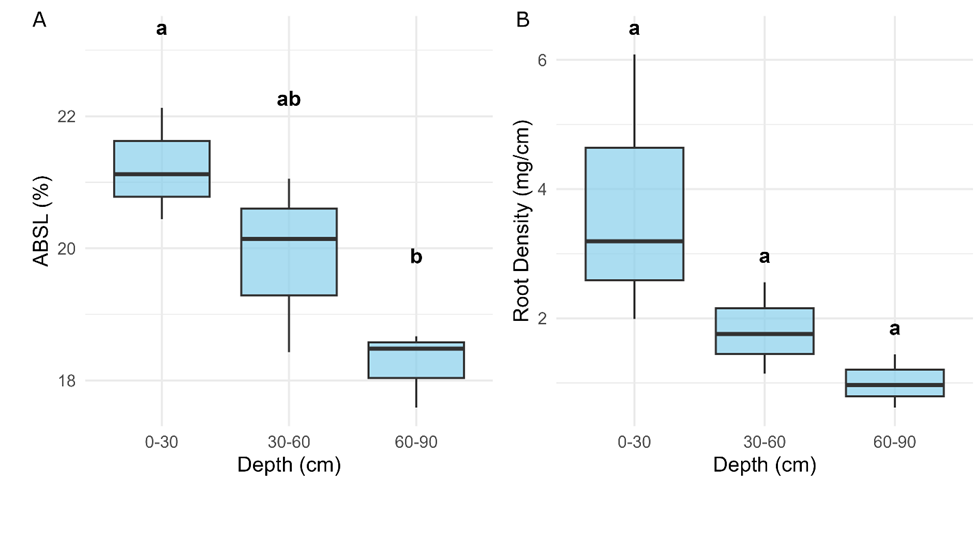

**Supplementary Figure S3.** Trends in lignin and root density variation for another lowland switchgrass genotype, AP13, are similar to the DVR genotype examined in depth. Roots were collected at 30 cm increments from 0-90 cm and analyzed for A) Acetyl bromide soluble lignin (ABSL) content and B) Primary root tissue density. Within a panel, significant differences between depths are indicated by different letters (ANOVA, Tukey HSD, P < 0.05, N = 3).

**Alt text_ Supplementary Figure S3**: Two boxplots showing changes in acetyl bromide soluble lignin content and root tissue density across four soil depth intervals (0-30, 30-60, and 60-90 cm) in the switchgrass accession AP13. In A) average ABSL is 21 % at 0-30 cm, 20% at 30-60 cm, and about 18.5% at 60-90 cm. Tukey’s HSD shows that 0-30 and 60-90 are significantly different from each other. In B) average root density is 3 mg/cm at 0-30 cm, 1.7 at 30-60 cm, and 1.0 at 60-90 cm. None of the distributions is statistically significantly different.
